# The shared ancestry between the C9orf72 hexanucleotide repeat expansion and intermediate-length alleles using haplotype sharing trees and HAPTK

**DOI:** 10.1101/2023.07.28.550820

**Authors:** Osma S. Rautila, Karri Kaivola, Harri Rautila, Laura Hokkanen, Jyrki Launes, Timo E. Strandberg, Hannu Laaksovirta, Johanna Palmio, Pentti J. Tienari

## Abstract

The C9orf72 hexanucleotide repeat expansion (HRE) is a common genetic cause of amyotrophic lateral sclerosis (ALS) and frontotemporal dementia (FTD). The inheritance is autosomal dominant, but a high proportion of cases are sporadic. One possible explanation is *de novo* expansions of unstable intermediate-length alleles (IAs). Using haplotype sharing trees (HST) with the novel haplotype analysis tool kit (HAPTK), we derived majority-based ancestral haplotypes of HRE carriers and discovered that IAs containing ≥18-20 repeats share large haplotypes in common with the HRE. Using HSTs of HRE and IA carriers, we demonstrate that the longer IA haplotypes are largely indistinguishable from HRE haplotypes. These analysis tools allow physical understanding of the haplotype blocks shared with the ancestral haplotype. Our results demonstrate that the haplotypes with longer IAs belong to the same pool of haplotypes as the HRE and suggest that longer IAs represent potential premutation alleles.

## Introduction

The GGGGCC hexanucleotide repeat expansion (HRE) in the first intron of the *C9orf72* gene is the most common cause of ALS and FTD in populations of European descent [1, 2, 3, 4]. Even though the inheritance is autosomal dominant, in many populations more than half of ALS cases with the C9orf72 HRE are sporadic [3, 4]. A founder haplotype has been reported in HRE carriers [5] and the same haplotype is commonly found in subjects with intermediate-length alleles (IAs, ≥7 repeats) indicating shared ancestry of the HRE and IAs [2, 5, 6, 7]. The HRE exhibits somatic instability (mosaicism) when repeat lengths have been compared between different tissues [8]. Whether the IAs also show instability is still unclear, although, in transfected cells DNA replication has been reported to stall at 20 GGGGCC-repeats [9]. Stalled DNA replication may recruit alternative, more error-prone replication methods, leading to both contractions and expansions of the repeat sequence [10].

Considering the hypothesis that IAs would be unstable premutations – as observed in other repeat expansion diseases [11] – the haplotypes with fewer repeats should be genetically more distant to HRE haplotypes. We set out to investigate the haplotype sharing between IAs and HRE carriers using haplotype sharing trees (HST) and majority-based ancestral haplotypes. To streamline the use of HST-based analyses in the future, we developed a haplotype analysis tool kit (HAPTK) written in the programming language Rust using HTSlib bindings from Rust-Bio [12, 13].

## Methods

### Cohorts and genotyping

To analyze HRE carriers, we used the Tampere ALS-FTD cohort (n=235 HRE carriers, 175 ALS, 55 FTD and 5 unclear cases), and the Helsinki ALS cohort (n=202 HRE carriers) [3]. For IA genotyping we selected 1474 samples from three Finnish population cohort studies (Supplements 1) [14, 15, 16] and 613 non-HRE ALS cases from the Helsinki ALS cohort.

The C9orf72 hexanucleotide repeat alleles were genotyped with repeat-primed PCR (RP-PCR), (Supplements 2). Because it may be difficult to distinguish between the HRE and a long repeat allele in RP-PCR, we confirmed alleles with ≥20 repeats and all putative expansions with over-the-repeat PCR [17, 18]. We also used the Amplidex C9orf72 method to ascertain a subset of our genotypes and observed high concordance with RP-PCR and AmplideX based genotypes (Supplements Table 1). Tampere ALS-FTD cohort was genotyped by over-the-repeat PCR and RP-PCR as part of diagnostic testing in the accredited Fimlab laboratories in Tampere.

Illumina Global Screening Array 24v2-3 was in used in SNP genotyping in all cohorts, except in the Helsinki ALS cohort, which was genotyped on FinnGen ThermoFisher Axiom custom array. After the quality control of each cohort (Supplements 3), we merged the cohorts and performed pre-phasing quality control on the merged dataset (Supplements 4) [19, 20]. Reliable HST construction requires the lowest possible switch-error and genotyping error rates. To minimize switch-errors, we used BEAGLE 5.3 [21, 22] and ran the phasing with 128 iterations. The variants were filtered for missingness (< 1%), and the missing genotypes were imputed during phasing. The Finnish SISU v3 population panel was used as the reference panel. After quality control and duplicate removal, Tampere ALS-FTD cohort consisted of 142 C9orf72 HRE ALS, 55 FTD and 5 unclear cases and the Helsinki ALS cohort of 187 C9orf72 HRE ALS cases.

### Uni- and bidirectional haplotype sharing trees (HST)

The construction of HSTs requires fully phased biallelic marker data which is read into a matrix of phased chromosomes (Fig. 1A and B, Supplements 6). Unidirectional HSTs are constructed separately down- and upstream of the starting marker. The algorithm moves along the matrix marker-by-marker, and if any contradictory alleles are found between the samples, the samples are branched into two nodes representing the diverged haplotypes. In contrast, the bidirectional HST is constructed by moving simultaneously in both directions, and when contradictory alleles are found on both sides, the samples are branched into two to four nodes. For both algorithms, the majority of samples are always assigned to the left-most node. After assigning the new leaf nodes, they are then recursively processed again to find contradictory alleles until all leaf nodes have only a single sample left, or the end of genotyping data is reached. The nodes contain information on the branching position and the indexes of the samples sharing the haplotype.

**Figure 1.**
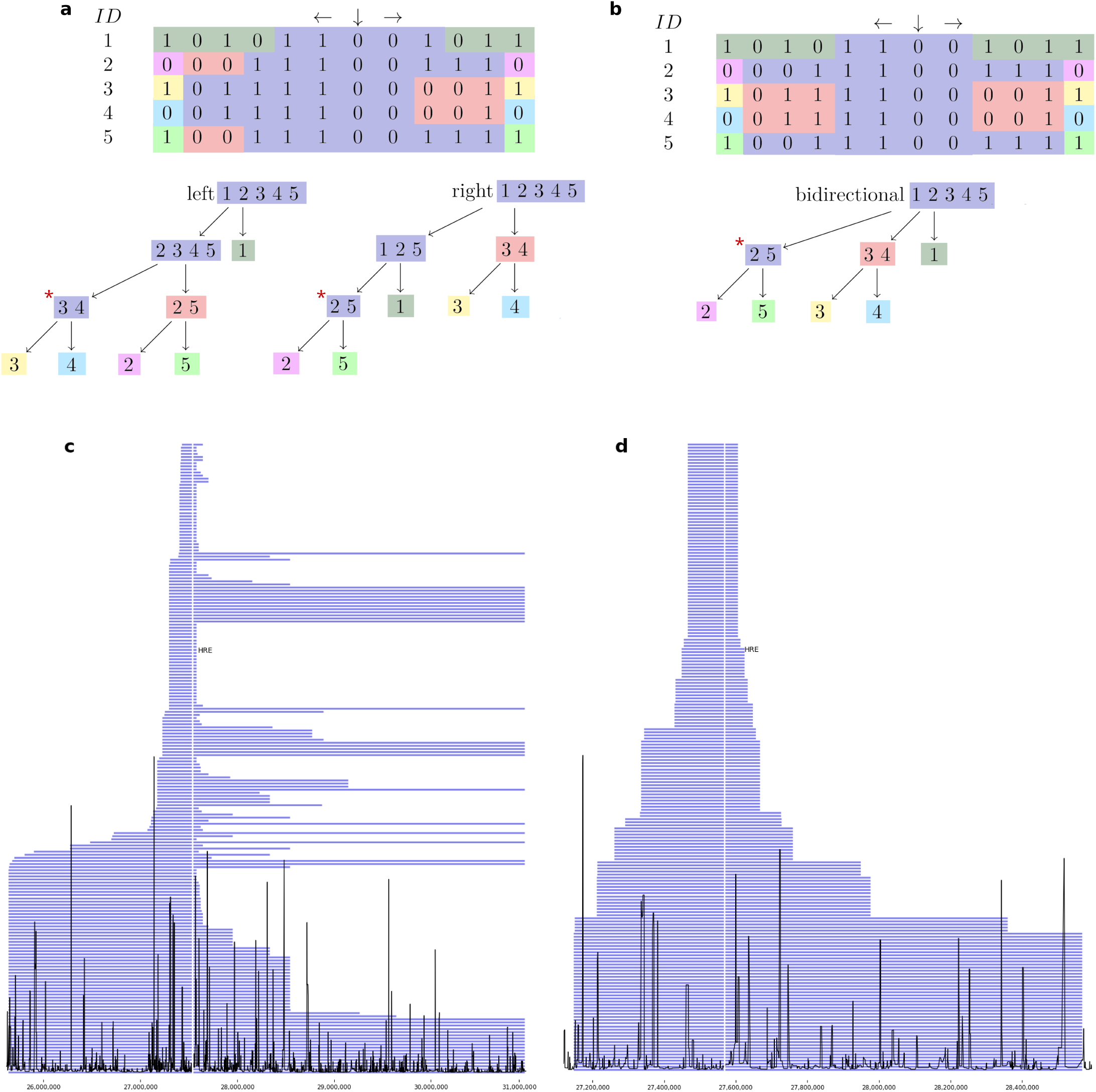
**a**. Illustration of the construction of unidirectional HST and **b**. bidirectional HST. Each column of the matrix is a biallelic SNP marker and each row represents a phased chromosome. The down-arrow denotes the starting marker of the algorithm. Given biallelic data, unidirectional HSTs are binary trees and bidirectional HSTs have three to four branches. * The node containing the majority-based ancestral haplotype. **c**. Segment lengths of the unidirectional HST-derived ancestral haplotype per sample after the selection for HRE haplotypes in the Tampere ALS-FTD cohort. Each row represents a single phased haplotype. Recombination frequencies are drawn in black and the vertical white line marks the HRE locus. **d**. Bidirectional HST-derived ancestral haplotype segment lengths after the selection for HRE haplotypes.

### The majority branch

The left-most branch of HSTs is called the *majority branch*. Following the majority branch we, in a way, walk back recombination events in time towards the ancestral haplotype. The haplotype in the last node on the majority branch, with a minimum of two samples, is the majority-based ancestral haplotype (Fig. 1A and 1B). With the unidirectional HSTs, the down- and upstream ancestral haplotypes are merged into a single ancestral haplotype (Fig. 1A).

### Switch-error and genotyping error evaluation

The majority-based approach effectively removes switch-errors from the ancestral haplotype, as for switch-errors to remain, they would need to be in the majority of samples. Therefore, to assess phasing quality, we compare all the haplotypes back against the ancestral haplotype. If long runs of shared ancestral segments are observed after the sample has separated from the majority, a switch-error or genotyping error has most likely caused it. To visualize this, we use the ancestral haplotype as a reference sequence and compare all the sample haplotypes to it (Fig. 2).

**Figure 2.**
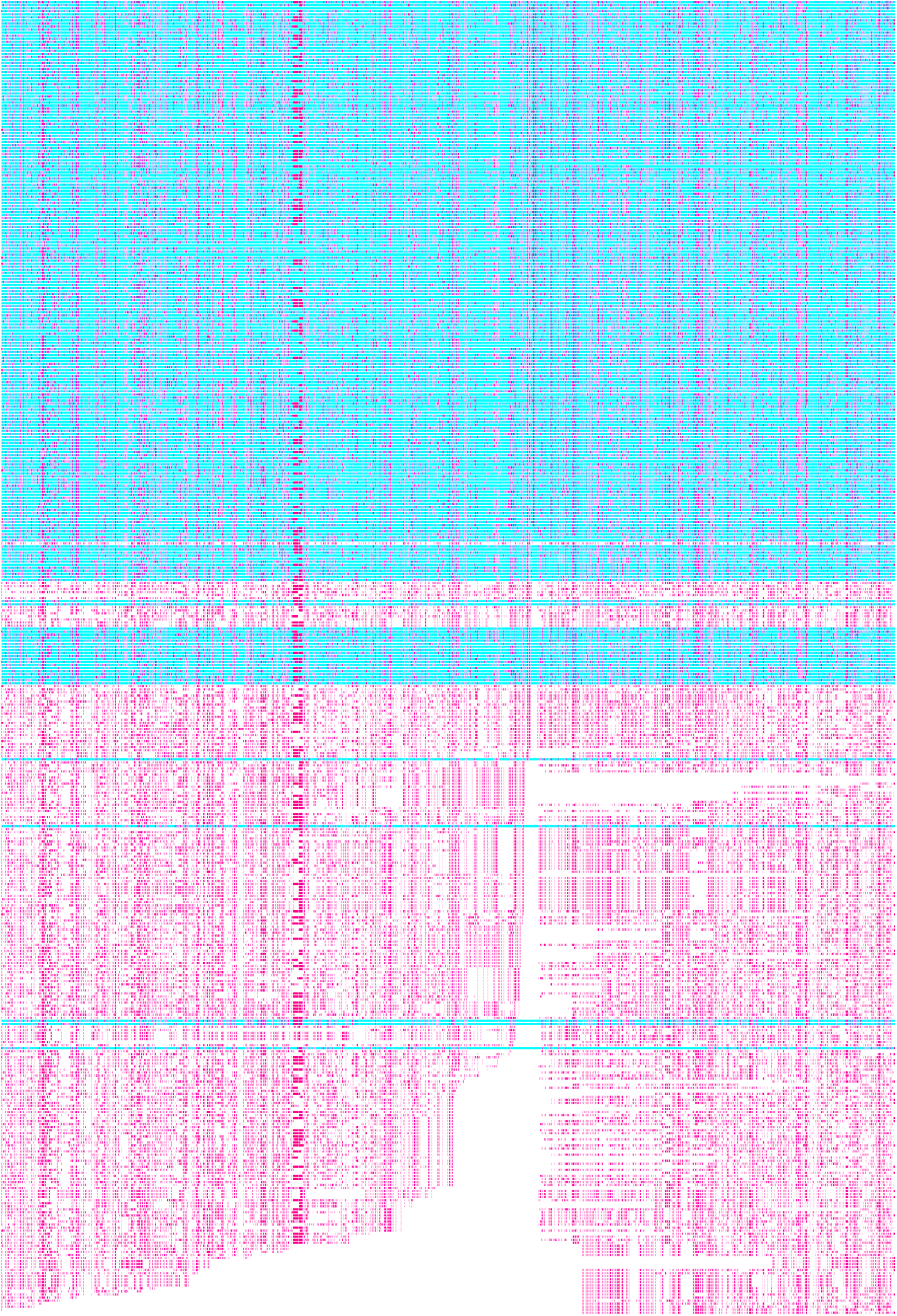
Phased haplotypes of the HRE samples aligned to the unidirectional HST-derived ancestral HRE haplotype as a reference. To visually separate each phased haplotype based on ancestral haplotype sharing, the reference alleles are colored in blue or white. We use blue for the haplotype sharing shorter segments per sample, and white for the one sharing longer segments. Alternative alleles are colored in pink. The selection of HRE haplotypes is done by discarding the haplotypes tagged in blue. As C9orf72 HRE inheritance is autosomal dominant, and no HRE homozygotes are present, the visual cut between blue and white phased haplotypes should be exact, with all the blue haplotypes on top. However, we found some IA haplotypes to share longer segments of the majority-based ancestral HRE haplotype than some HRE haplotypes themselves. For example, the lowest three haplotypes tagged in blue carry both the HRE and ≥20 repeat IAs. Above the last blue line, a long segment of downstream (left side) ancestral sequence is cut off indicating a possible switch-error.

### Selecting HRE haplotypes

As no SNPs tag only the HRE [18], we select the HRE haplotypes by comparing the lengths each phased haplotype shares with the ancestral HRE haplotype. For each sample, we remove the haplotype that shares a shorter segment of the ancestral HRE haplotype (tagged in blue in Fig 2). After selecting only the longer segments, we reconstruct the HST.

### MRCA estimation with the Gamma method

The lengths of shared segments with the ancestral HRE haplotype can be used to infer how much recombination has occurred in the cohort. This is the basis of MRCA estimation with the Gamma method [23]. We use the unidirectional HST and register the sharing with the ancestral HRE haplotype for each sample. We then translate the distance between the breakpoint from the ancestral haplotype and the HRE locus to centimorgans using Beagle genetic maps. To estimate the MRCA, we ported the original algorithm without chance-sharing correction [23] from R into our application.

### Analysis of the HRE and IA ancestries

The IAs were categorized into predefined groups of 2-6, 7-9, 10-14, 15-19, and 20-45 repeats [18]. The cut-off point of 7 repeats is commonly used to define the smallest IA, the cut-off 10 is based on a peak in allele frequency, the cut-off 15 is based on a drop in allele frequency and the cut-off 20 is based on a tagging SNP [18]. To estimate a possible threshold effect, we made an additional analysis with discrete IAs.

## Results

We constructed the HSTs and derived the core and ancestral HRE haplotypes using a starting base pair position 27,582,313 (GRCh38) which was the closest marker upstream of the C9orf72 HRE. In the Tampere ALS-FTD cohort we found a 13-SNP 114 kb core haplotype, shared by all HRE carriers (Supplements table 2). We visualized the individual shared ancestral HRE haplotype segment lengths in Fig. 1C and 1D. In the Helsinki ALS cohort, we found a 10-SNP 89 kb core HRE haplotype (Supplements table 3).

Next, we visualized the unidirectional (Fig. 3, Supplements Fig 4.) and bidirectional (Supplements Fig 2 and 5) HSTs. Based on relatively early upstream recombination, the Tampere ALS-FTD cohort seemed to form two subtrees. We then tagged the FTD cases in the bidirectional HST to analyse whether FTD haplotypes would differ from ALS haplotypes, however, the FTD cases did not form any clear subtrees indicating that the haplotype composition is not a major determinant, whether the subject develops ALS or FTD (Supplements Fig. 3).

**Figure 3.**
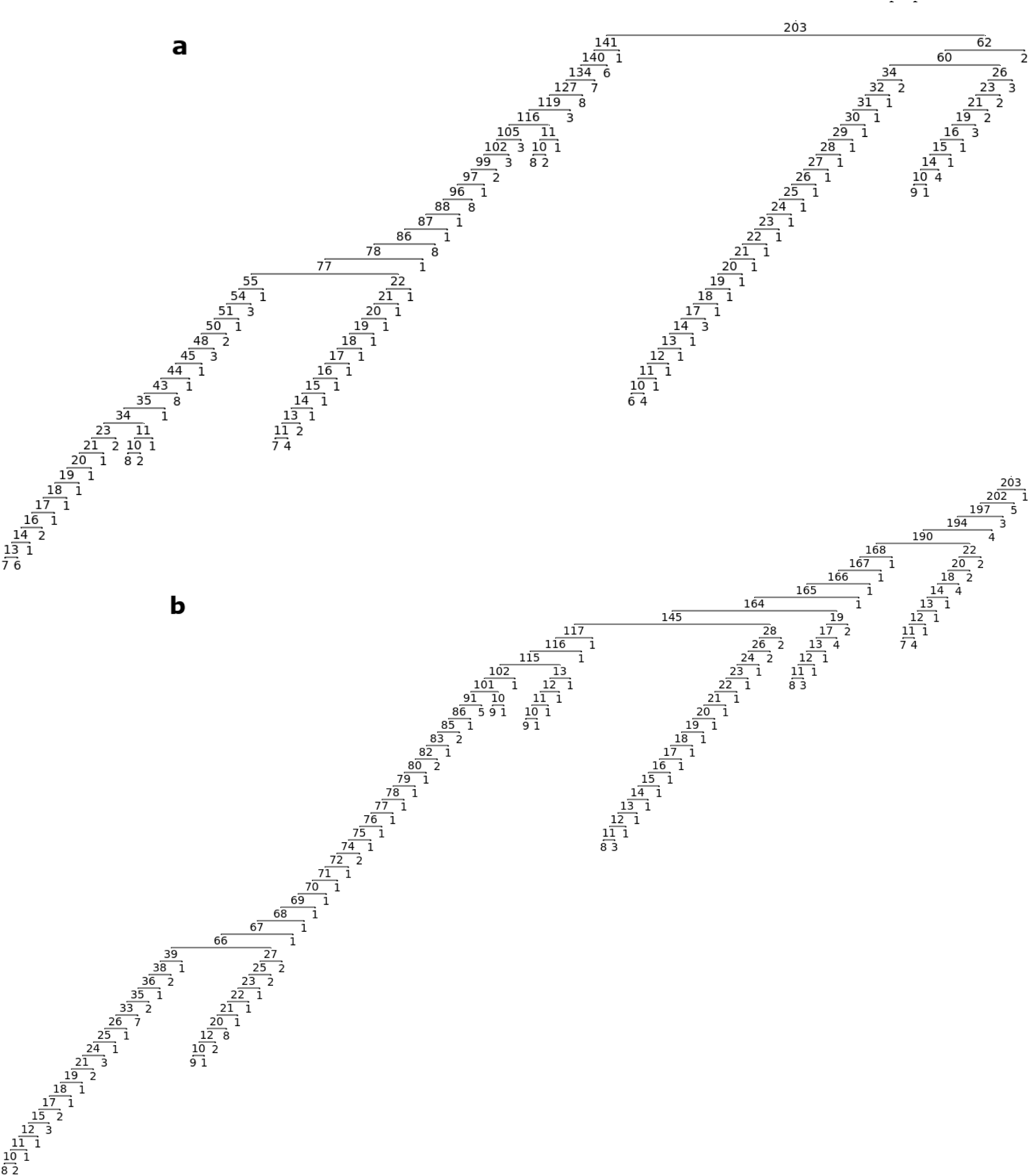
**a**. The right side (upstream) unidirectional HST. **b**. The left side (downstream) unidirectional HST. For visualization purposes, the branching is stopped after nodes have less than 10 samples remaining.

To assess the quality of the phasing and the integrity of the HSTs, we compared all the haplotypes against the unidirectional HST-derived ancestral HRE haplotype. The comparison graph was sorted by the amount of downstream (left side) haplotype sharing between the sample and the ancestral HRE haplotype (Fig. 2).

Based on the comparison graph, switch-error rate was visually determined to be low, and we chose not to try to correct the few potential switch-errors remaining in the data. The selection of HRE haplotypes based on ancestral HRE haplotype sharing was deemed adequate (Fig. 2).

Then, we compared predefined groups of IAs to the unidirectional HST-derived ancestral HRE haplotype and found most haplotype sharing in the ≥20 repeat IA category in both cohorts. Haplotype sharing strongly diminished already in the 15-19 repeat IA category and continued to decrease towards the 2-6 repeat category (Fig. 4A). To detect a possible threshold effect, we analyzed unidirectional HST-derived ancestral haplotype sharing for all haplotypes with ≥10 repeat discrete IAs. A threshold effect was seen in IA haplotypes starting at 18 repeats (Fig. 4I).

**Figure 4.**
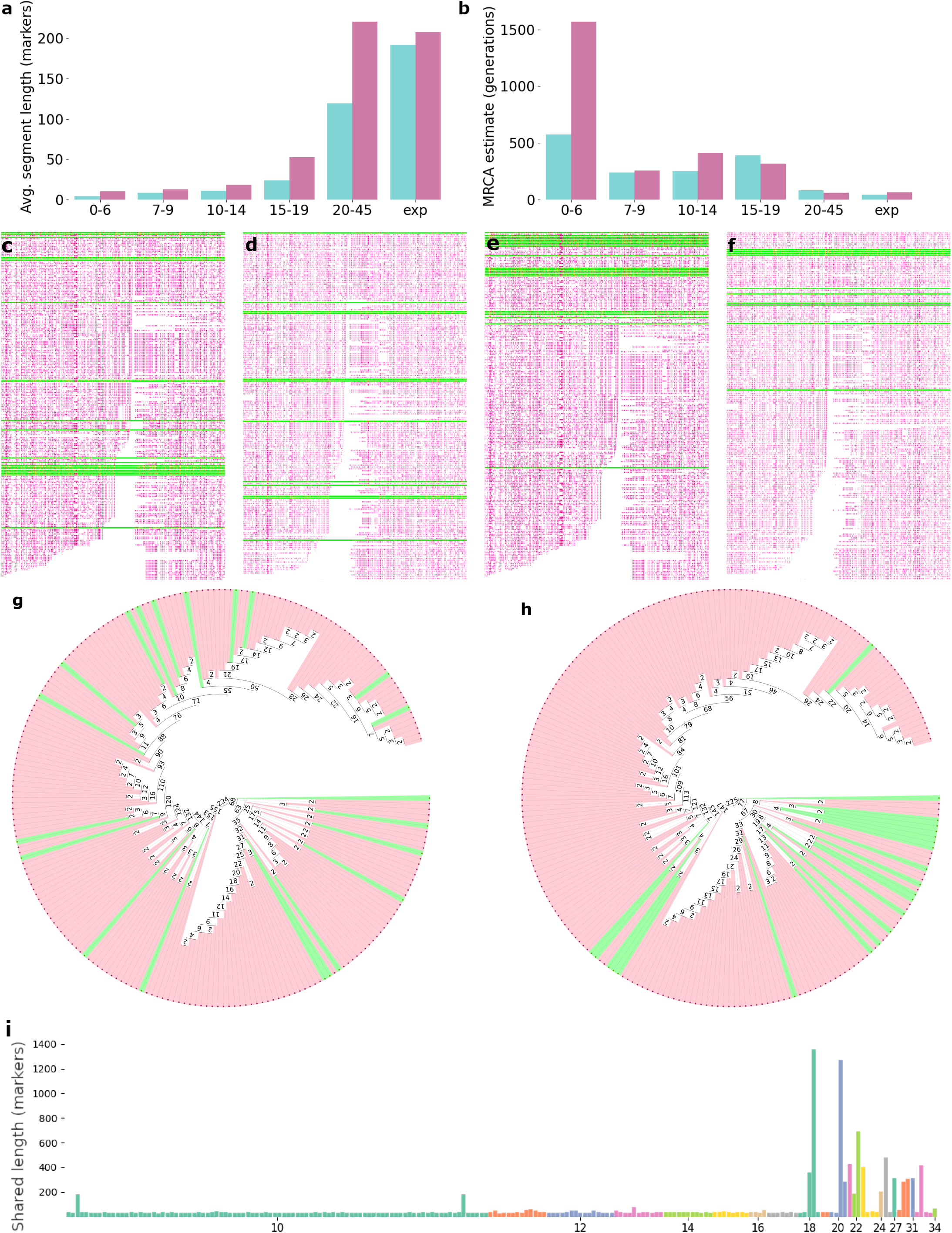
**a**. Average ancestral HRE haplotype segment length in markers per IA group. In the Helsinki ALS cohort in the ≥20 IA group nine out of twelve were carriers of two longer ≥7 IAs, raising the average segment sharing over the HRE group itself. **b**. MRCA estimates in generations using the Gamma method with correlated genealogy. **c**. Tampere ALS-FTD cohort HRE carriers mixed with ≥20 repeat IAs and compared against the unidirectional HST-derived ancestral haplotype. IA haplotypes are tagged in green. **d**. Helsinki ALS cohort HRE carriers mixed with ≥20 repeat IAs. **e**. Tampere ALS-FTD cohort HRE carriers mixed with 15-19 repeat IAs. **f**. Helsinki ALS cohort HRE carriers mixed with 15-19 repeat IAs. **g**. A bidirectional HST of Tampere ALS-FTD HRE carriers mixed with ≥20 IAs and ≥20 IAs tagged in green and **h**. the HRE carriers mixed with ≥15 IAs and ≥15 IAs tagged in green. **i**. Ancestral HRE haplotype segment lengths (y-axis) in ≥10 repeat groups for both phased chromosomes of all the Tampere ALS-FTD and population control samples. For visualization purposes, all haplotypes sharing less ancestral sequence than the core HRE haplotype consisting of 11 SNPs were filtered out on the x-axis.

Next, we compared all the haplotypes to the unidirectional HST-derived ancestral HRE haplotype and tagged the IAs in green. The ≥20 repeat IAs were seen to distribute relatively evenly along the graph in both cohorts (Fig. 4C and D). On the other hand, most of the 15-19 repeat IAs diverged from the HRE majority quite early (Fig. 4E and F). To detect possible division into subtrees, we constructed a bidirectional HST of the ≥20 repeat IAs mixed with HRE alleles and tagged the IA carriers in red. We found that the ≥20 repeat IAs were spread across the HSTs and did not form subtrees within the HRE carriers (Fig. 4G, Supplements Fig. 6).

Finally, we estimated the MRCA with the Gamma method under correlated genealogy and found that in the shorter groups MRCA was high, but in the ≥20 intermediate alleles, it was close to the HRE in both cohorts (Fig. 4B). The MRCAs of the HRE carriers were ca. 54.6 (9.7 – 98.1, 95% CI, Tampere ALS-FTD cohort) and ca. 65.3 (14.8 – 114.7, 95% CI, Helsinki ALS cohort) generations ago.

## Discussion

When comparing the ancestral HRE haplotype sharing in different length IAs, we found significantly increased haplotype sharing starting from 18-20 repeats, however the number of observations did not allow precise estimation of the threshold. These findings demonstrate similarly shared ancestry between carriers of the longer IAs and the HRE. More than half of ALS patients with the C9orf72 HRE are sporadic [3, 4] and one hypothesis for this observation is parental pre-mutation, i.e., an unstable IA [7, 18, 24, 25]. The population frequency of ≥20 repeat alleles is around 1.7% [17], and the frequency of the HRE in older population is in the range of 0.2% [24]. Given the high degree of sharing between HRE and longer IAs, our data supports the concept that the expanded alleles are formed from the longer IAs. However, whether some IAs are generated from contracted expansions cannot be ruled out. These findings align with studies showing that at 20 GGGGCC repeats, the DNA replication system becomes more error-prone [9].

This study has certain limitations. First, some IA alleles share long ancestral HRE haplotypes, and as a few HRE carriers also carry IAs, the selection of the longer ancestral HRE haplotype segment does not mean it is certain that the HRE allele is selected. For example, we found three subjects with both the HRE and ≥20 repeat IAs, but when tested, the removal of these samples did not affect the ancestral haplotype. Second, a few possible switch-errors were seen in the comparison graphs, and corrections were not attempted on them. We believe the effects of these few switch errors are negligible.

Regarding ancestral haplotype sharing, the uni- and bidirectional HST-derived analyses give similar results. The benefit of unidirectional HST is that recombination events are decoupled on both sides of the target, however bidirectional HSTs are beneficial when a single tree simplifies downstream analyses. The future use for bidirectional HST include haplotype block-based genome-wide association studies by stepwise genome-wide construction of HSTs and then minimizing each tree for a minimum node value (in associations studies, the p-value from comparing node samples to the rest of the samples). HST can also be used to study haplotype effects on quantitative variables. Although, the analyses on these cohort sizes were run within minutes on consumer grade hardware, and memory usage was negligible, several optimizations remain to be done to allow for efficient use with biobank scale data.

## Code availability

HAPTK and all the scripts used in this article are available at https://github.com/xosxos/haptk

## Acknowledgements

The data used for the research was imputed with the THL Biobank’s SISU v3 Imputation reference panel obtained from THL Biobank. We thank all study participants for their generous participation in the FINRISK, Health 2000, and Migraine Family studies. We also thank the Sequencing Informatics Team, FIMM Human Genomics, University of Helsinki for the work done in preparation of the reference panel data. This study was funded by the Finnish Cultural Foundation, Päivikki and Sakari Sohlberg Foundation, Paulo Foundation, the Sigrid Juselius Foundation, Maire Taponen Foundation, the Finnish Brain Foundation, the Finnish Medical Foundation, the Biomedicum Helsinki Foundation, Helsinki University Hospital grants, ALS tuttu ry and the Finnish Academy (318868).

