## supplements for "The shared ancestry between the C9orf72 hexanucleotide repeat expansion and intermediate-length alleles using haplotype sharing trees and HAPTK"

#### 1. Control cohorts

The Helsinki Businessmen study (HBS) included men with high socio-economic status who were born in 1919–1934 and in 2002–2003, 672 individuals who lived at home were randomly selected for analyses (DNA was available in 666).

The DEBATE study was originally a random sample of 4800 individuals from Helsinki out of whom 400 home-living individuals with stable cardiovascular disease were randomly selected for further studies in 2000 (DNA was available for 375). We excluded one individual diagnosed with ALS from these cohorts, which decreased the number of controls from the original 3142 to 3141.

The PLASTICITY cohort is an ongoing long-term follow-up of originally 1196 individuals born in 1971–74 in the Helsinki metropolitan area who had at least one predefined pre- or perinatal risk factor (e.g. low birth weight or Apgar score). 509 subjects were seen at age 40 (DNA available in 433).

#### 2. Genotyping C9orf72 hexanucleotide repeats

We extracted DNA using standard methods from peripheral blood leukocytes or saliva (PLASTICITY cohort). For all samples we first assessed repeat lengths using repeat-primed PCR (RP-PCR) followed by capillary electrophoresis and the results were visualized using the GeneMapper software v6 (ThermoFisher). Then, for samples with a putative expansion (including all samples with $\geq 20$ repeats) or unclear zygosity, we performed over-the-repeat PCR. Samples that showed the typical saw tooth pattern in RP-PCR and did not produce longer amplicon in over-the-repeat PCR were categorized as expansions. The longest non-expanded (amplifiable) discrete allele we could detect was 45 repeats, and we used it as the expansion threshold.

For quality control, we examined the genotyping concordance between AmplideX C9orf72 kit and our RP-PCR.

**Supplements table 1. Genotype concordance between AmplideX C9orf72 kit and RP-PCR**

| Sample | Amplidex genotype (manually checked genotypes when genotype missing or mismatching) | RP-PCR genotype |
|---|---|---|
| A14LIALS101 | 2, >145 | 2/exp |
| A14LIALS15 | 8, 8 (8/10) | 8/10 |
| A14LIALS169 | 2, 2 (2/exp) | 2/exp |
| A14LIALS187 | 8, 35 | 8/36 |
| A14LIALS236 | 2, 19 | 2/19 |
| A14LIALS26 | 8, 22 | 8/22 |
| A14LIALS272 | 2, 36 | 2/37 |
| A14LIALS279 | 2, 11 | 2/11 |
| A14LIALS283 | 2, 19 | 2/19 |
| A14LIALS285 | 2, 12 (2/12) | 12/12* |

| | | |
|---|---|---|
| A14LIALS300 | 2, >145 | 2/exp |
| A14LIALS301 | 2, >145 | 2/exp |
| A14LIALS303 | 2, >145 | 2/exp |
| A14LIALS312 | 2, 2 (2/exp) | 2/exp |
| A14LIALS391 | 5, 15 | 5/15 |
| A14LIALS405 | 2, 14 | 2/14 |
| A14LIALS45 | (2/36) | 2/37 |
| A14LIALS452 | 8, 22, 41 | 8/22* |
| A14LIALS58 | 5, >145 | 5/exp |
| A14LIALS66 | (2/exp) | 2/exp |
| N19ALS172 | 2, >145 | 2/exp |
| N19ALS176 | 5, >145 | 5/exp |
| N19ALS187 | 2, >145 | 2/exp |
| N19ALS20 | 7, 16, 34 (7/16) | 7/16 |
| N19ALS225 | 2, >145 | 2/exp |
| N19ALS254 | (2/exp) | 2/exp |
| N19ALS326 | 2, >145 | 2/exp |
| N19ALS337 | 10, 18 | 10/18 |
| N19ALS35 | 2, 22 | 2/22 |
| N19ALS43 | 5, 23 | 5/23 |
| N19ALS53 | 8, 10 | 8/10 |
| S14F13ALS19 | 7, 24 | 7/24 |
| S14F13ALS37 | 2, 2, >145 (2/145) | 2/exp |
| S14F13ALS45 | 2, >145 | 2/exp |
| S14F13ALS54 | 8, >145 | 8/exp |
| S14F13ALS62 | 8, 8 (8/13) | 8/13 |
| S14F13ALS77 | 2, 45 (2/42-48) | 2/exp |
| S14F13ALS79 | 5, >145 | 5/exp |
| S14F13ALS8 | 8, >145 | 8/exp |
| S14F13ALS80 | 2, >145 | 2/exp |

*Genotype confirmed with over-the-repeat PCR

#### 3. SNP data quality control

Genotyping data quality control was performed separately on samples processed with different arrays. First, we removed duplicated samples and related samples (proportion IBD > 0.1875), samples with discordant sex information, outlying heterozygosity rate (>3SD) and > 5% missing genotype rate. In per-variant quality-control, we included variants that had a genotyping rate of > 95%, variants in HWE (p>0.000001) and minor allele frequency >= 0.01. To harmonize genotyping array data, we included only biallelic non-palindromic SNPs and set allele coding to match the reference genome so that A1 allele was always the alternative allele regardless of minor allele frequency.

#### 4. Phasing quality control and imputation

First, we excluded markers with discrepant AF to the SISU v3 reference panel or not present in the reference panel. We performed phasing with *ne* set to 20 0000 and number of iterations set to 128.

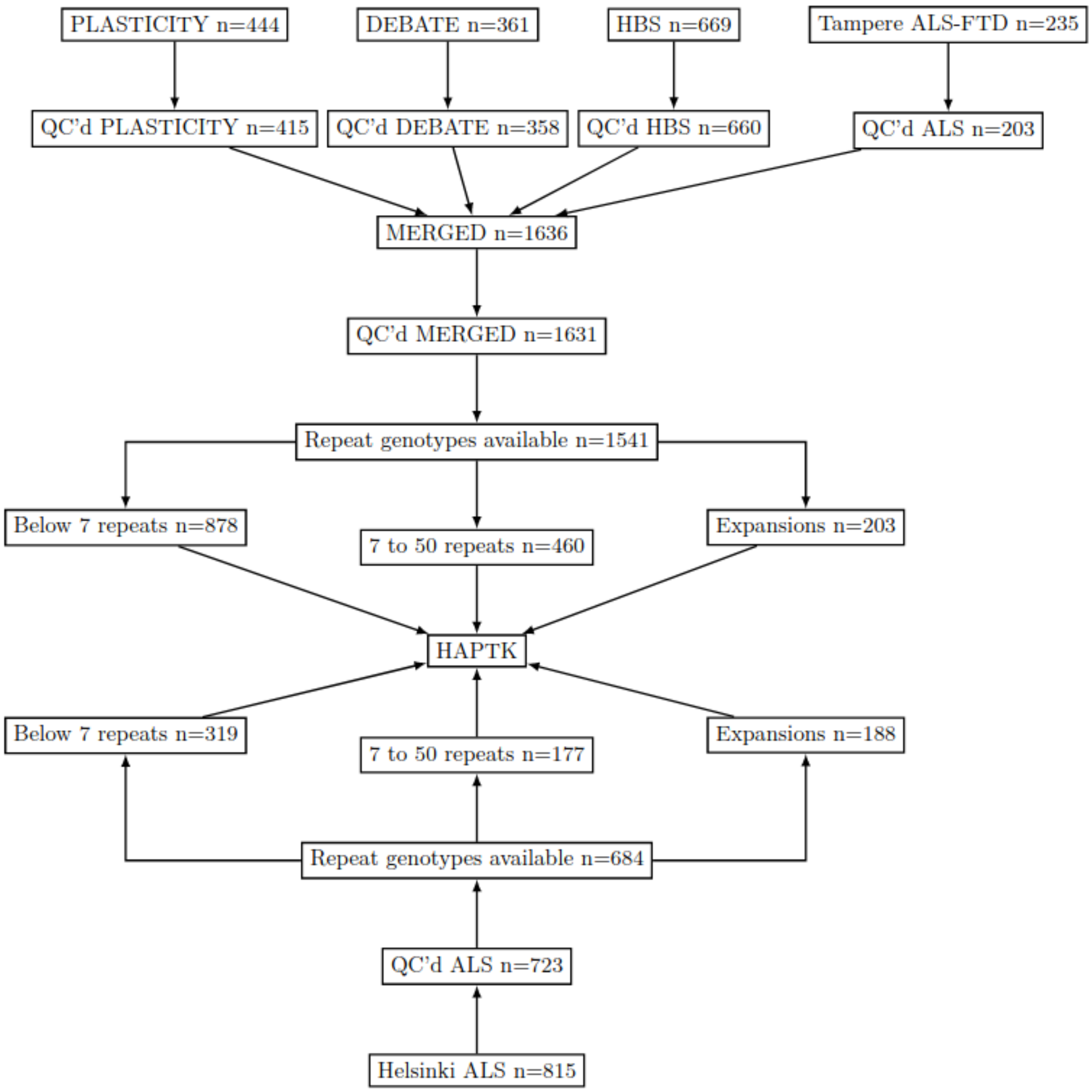


**Supplements figure 1. Samples used in the study. Illumina GSA v2-3 chip cohorts on the top and the Affymetrix Axiom Custom SNP array Helsinki ALS samples on the bottom.**

#### 5. Unidirectional haplotype sharing tree algorithm

The genotype matrix consists of biallelic phased data without missing genotypes. In the sample-variant matrix, each sample has one row of genotypes for each of the two haploid genotypes.

Outline of the algorithm:
1. Select a wanted starting marker (usually the locus of the studied variant) from the genotype matrix by a coordinate, if the given coordinate has no designated variant, select the one immediately to the right

2. Travel marker by marker to both sides of the target separately.

3. When a contradictory genotype between the haploid genotypes is found, assign samples into two new nodes based on the genotypes.  For biallelic data two possible genotypes are present [0], [1]
Z
6. Continue the traversal by selecting only leaf nodes and perform the search for the next contradictory genotype, but only in the context of samples present in the leaf node.

7. Continue until all leaf nodes have only a single sample left or sequencing data runs out

#### 6. Bidirectional haplotype sharing tree algorithm

The genotype matrix consists of biallelic phased data without missing genotypes. In the sample-variant matrix, each sample has one row of genotypes for each of the two haploid genotypes.

1. Select a wanted starting marker (usually the locus of the studied variant) from the genotype matrix by a coordinate, if the given coordinate has no designated variant, select the one immediately to the right

2. Travel to the right until a contradictory genotype is found between samples.

3. Travel to the left until a contradictory genotype is found between samples.

4. When both sides present a contradictory genotype, assign samples to four possible buckets: [0,1], [1,0] [0,0] [1,1]. For each bucket, add a new node into the tree.

6. Continue the traversal by selecting only leaf nodes and perform the search for the next contradictory genotypes, but only in the context of samples present in the leaf node.

7. Continue until all leaf nodes have only a single sample left or sequencing data runs out

#### 7. Additional tables

| pos | ref | alt | genotype |
|---|---|---|---|
| 27474216 | T | C | 0 |
| 27477876 | C | T | 1 |
| 27478711 | T | C | 1 |
| 27484498 | A | G | 0 |
| 27502988 | C | A | 0 |
| 27508689 | T | C | 1 |
| 27510494 | C | T | 1 |
| 27513838 | C | T | 0 |
| 27546892 | G | A | 0 |
| 27553878 | T | C | 0 |
| 27556782 | G | A | 0 |
| 27557532 | T | C | 0 |
| 27573534 | GGGGCC | HRE | 1 |
| 27587790 | T | C | 0 |

**Supplements table 2: Tampere ALS-FTD cohort. HRE carrier core haplotype aligned to GRCh38 (Illumina Global Screening Array 24v3)**

| pos | ref | alt | genotype |
|---|---|---|---|
| 27496247 | T | C | 0 |
| 27498553 | G | T | 0 |
| 27511845 | G | A | 0 |
| 27549487 | G | T | 0 |
| 27556782 | G | A | 0 |
| 27557532 | T | C | 0 |
| 27564340 | T | C | 0 |
| 27570053 | A | G | 0 |
| 27573534 | GGGGCC | HRE | 1 |
| 27557532 | T | C | 1 |
| 27573534 | T | C | 0 |

**Supplements table 3: Helsinki ALS cohort. HRE carrier core haplotype aligned to GRCh38 (Affymetrix Axiom Custom SNP Array)**

### 8. Additional figures

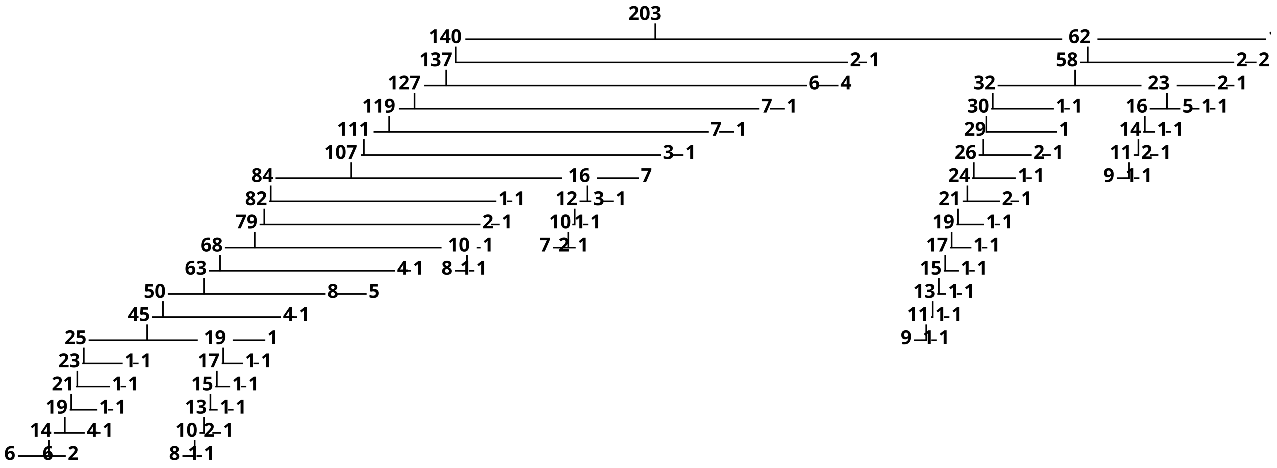

**Supplements figure 2: Bidirectional HST of HRE cases in the Tampere ALS-FTD cohort genotyped on the Illumina GSA v3 SNP array. For visualization purposes the branching is cut after 10 samples.**

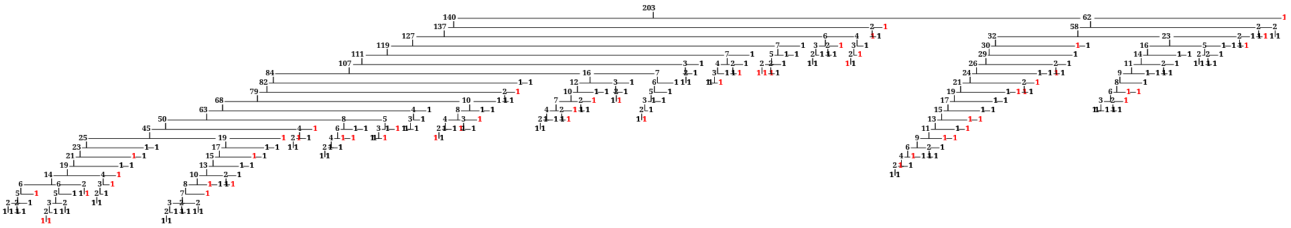

**Supplements figure 3: Bidirectional HST of HRE cases with FTD cases tagged in red in the Tampere ALS-FTD cohort.**

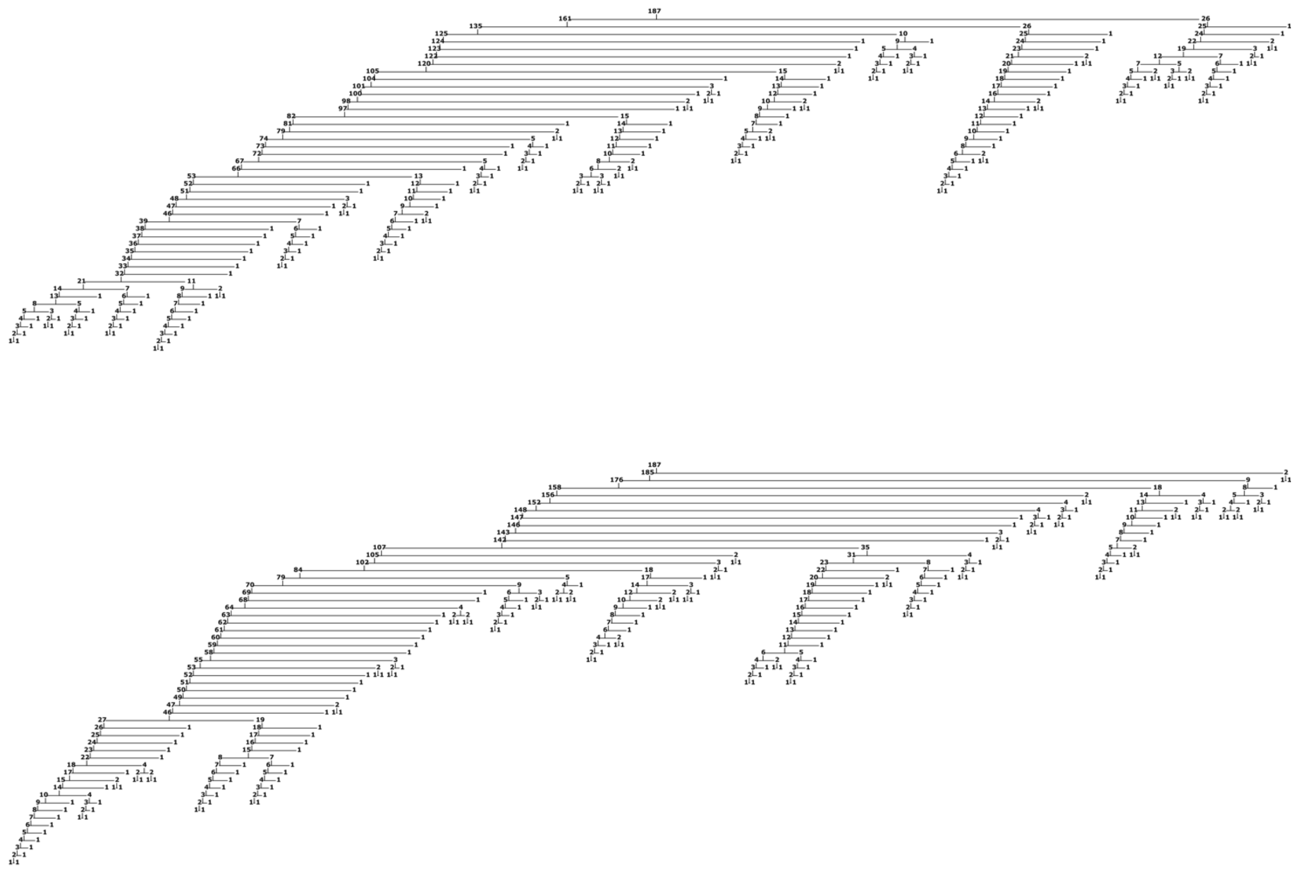

**Supplements figure 4: Unidirectional HST of HRE cases in the Helsinki ALS cohort genotyped on the Affymetrix Axiom SNP array. Right side on the top. Left side on the bottom.**

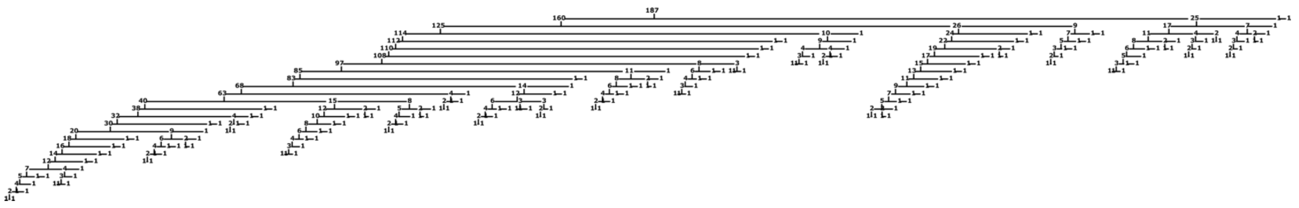

**Supplements figure 5: Bidirectional HST of HRE cases in the Helsinki ALS cohort genotyped on the Affymetrix Axiom SNP array.**

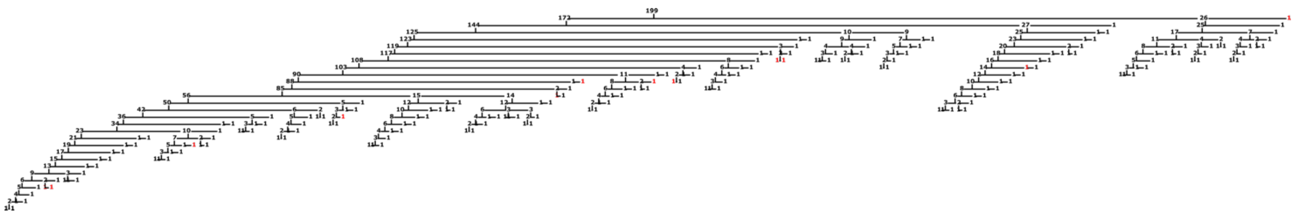

**Supplements figure 6: Bidirectional HST of HRE and ≥20 repeat IAs. The ≥20 repeat IAs are tagged in red.**
